# Genetic Nature or Genetic Nurture? Quantifying Bias in Analyses Using Polygenic Scores

**DOI:** 10.1101/524850

**Authors:** Sam Trejo, Benjamin W. Domingue

**Affiliations:** Graduate School of Education, Stanford University

**Author notes:** Send correspondence to.

## Abstract

Summary statistics from a genome-wide association study (GWAS) can be used to generate a polygenic score (PGS). For complex, behavioral traits, the correlation between an individual’s PGS and their phenotype may contain bias alongside the causal effect of the individual’s genes (due to geographic, ancestral, and/or socioeconomic confounding). We formalize the recent introduction of a different source of bias in regression models using PGSs: the effects of parental genes on offspring outcomes, also known as genetic nurture. GWAS do not discriminate between the various pathways through which genes influence outcomes, meaning existing PGSs capture both direct genetic effects and genetic nurture effects. We construct a theoretical model for genetic effects and show that, unlike other sources of bias in PGSs, the presence of genetic nurture biases PGS coefficients from both naïve OLS (between-family) and family fixed effects (within-family) regressions. This bias is in opposite directions; while naïve OLS estimates are biased upwards, family fixed effects estimates are biased downwards. We quantify this bias for a given trait using two novel parameters that we identify and discuss: (1) the genetic correlation between the direct and nurture effects and (2) the ratio of the SNP heritabilities for the direct and nurture effects.

## 1. Introduction

### 1a. Genomics & the Social Sciences

Spurred by the plummeting cost of DNA sequencing and technological developments in processing large amounts of genetic data, researchers have made great strides in connecting genes to biological and social outcomes in a replicable manner. The key tool is the genome-wide association study (GWAS); a GWAS uses genotype and phenotype data from many individuals to probe the relationship between a given trait and thousands of regions of the genome (Pearson and Manolio 2008). GWAS are conducted on a wide variety of outcomes, ranging from proximal, biological phenotypes, such as blood pressure (Giri et al. 2019) and height (Yengo, Sidorenko, et al. 2018), to distal, behavioral phenotypes, such as depression (Hyde et al. 2016; Okbay et al. 2016) and educational attainment (Lee et al. 2018).

Findings from GWAS are often used to generate a predictor—a polygenic score (PGS)— meant to summarize an individual’s genetic predisposition for a given trait. PGSs offer great promise to social scientists interested in incorporating genes into biosocial models of human behavior (Belsky and Israel 2014). In the short term, PGSs may be used as control variables in studies of environmental effects (Rietveld et al. 2013), used in gene-environment interaction studies to probe whether genetic effects are environmentally contingent (Trejo et al. 2018; Barcellos, Carvalho, and Turley 2018), and used to better understand how genetic factors influence developmental processes (Belsky et al. 2016; Belsky et al. 2013). In the long run, PGSs might be used to identify those who would benefit most from early medical or educational interventions, i.e., for a developmental disorder like dyslexia (Martschenko, Trejo, and Domingue 2019).

### 1b. The Problem of Confounding

A point of emphasis is that the same technique, GWAS, is being used to map the genetic architecture of a diverse set of phenotypes. It is not obvious that the methodology used to identify the underlying genetics of proximal, biological phenotypes can be deployed without side effect to interrogate the genetics of complex, socially contextualized phenotypes. Especially in the case of traits like depression and educational attainment, it is critical that existing GWAS results be interpreted cautiously (Martschenko, Trejo, and Domingue 2019); while PGSs have been shown to predict complex phenotypes, the relationship between an individual’s PGS captures a broad range of information and associations with downstream outcomes and therefore cannot be readily interpreted as the causal effect of genes. An individual’s genome contains fine-grain information about their place in the intricate structure of a population (Hamer and Sirota 2000; Novembre et al. 2008), meaning that GWASs for complex traits may simply identify genes related to confounding environmental variables such as ancestry, geography, or socioeconomic status.

Recent work in human studies has begun to elucidate a novel source of confounding: social genetic effects (Domingue and Belsky 2017). Social genetic effects, also known as indirect genetic effects, are defined as the influence of one organism’s genotype on a different organism’s phenotype. The idea of social genetic effects originated in evolutionary theory (Moore, Brodie, and Wolf 1997; Wolf et al. 1998), and social genetic effects have been observed in animal populations (Petfield et al. 2005; Bergsma et al. 2008; Canario, Lundeheim, and Bijma 2017; Baud et al. 2018). Social science is now beginning to study such effects in human populations; examples include among social peers (Sotoudeh, Conley, and Harris 2017; Domingue et al. 2018), sibling pairs (Cawley et al. 2017; Kong et al. 2018), and parents and their children (Bates et al. 2018; Kong et al. 2018; Wertz et al. 2018). The existence of within-family social genetic effects complicates attempts to derive causal estimates from GWAS.

For recent breakthroughs in the genetic architecture of complex traits to provide novel value to researchers in the biomedical and social sciences, the relationships discovered in a GWAS must mostly reflect causal relationships between an individual’s genes and their phenotype. If, for example, the genes identified for a complex trait predict it only through spurious correlation, PGSs will provide little use towards broadening our understanding of genetic and environmental influences. Thus, validating PGSs within-families is vitally important for sifting out causation from correlation among the genetics identified in GWASs of complex traits (Rietveld et al. 2014; Domingue et al. 2015; Lee et al. 2018; Belsky et al. 2018). Environmental differences are muted between siblings and, conditional on parental genotype, child genotype is randomly assigned through a process known as genetic recombination (Conley and Fletcher 2017). This makes family fixed effect regression models that compare genetic differences in siblings to phenotypic differences in siblings the gold standard for testing and understanding whether genetic differences are causally related to downstream outcomes. Within-family research designs, however, are not without their own complications. Genetic nurture may lead to bias in estimates derived from within-family studies, though the extent of this bias has not yet been explored.

### 1c. Accounting for Genetic Nurture

In this paper, we describe how genetic nurture influences PGS construction and leads to bias within-family and between-family regression analyses using PGSs. We construct a theoretical model for additive genetic effects and show that, unlike other sources of bias in PGSs, the presence of genetic nurture can bias PGS coefficients from <u>both</u> naïve OLS (between-family) regressions and family fixed effects (within-family) regressions. We quantify the magnitude of this bias for a given trait using two novel parameters: (1) the genetic correlation between the direct and nurture effects and (2) the ratio of the SNP heritabilities for the direct and nurture effects. Bias is in opposite directions; whereas naïve OLS estimates are biased upwards, family fixed effects estimates are biased downwards. These findings highlight a shortcoming of existing PGSs and have important implications for the use and interpretation of research designs using PGSs for traits where genetic nurture is a relevant causal pathway.

The paper will proceed as follows. In Section 2, we motivate our theoretical model using empirical data. We introduce our theoretical model in Section 3 and then demonstrate its implications for GWAS and regression models using PGSs in Section 4. In Section 5, we discuss the insights gleaned from our theoretical model.

## 2. Empirical Motivation

### 2a. Empirical Model

We motivate our theoretical framework by first considering the empirical specifications used in recent work (Domingue et al. 2015; Belsky et al. 2018; Lee et al. 2018). Consider the following two models relating an individual’s PGS constructed from recent GWAS results 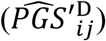 to their outcome (*Y*_*ij*_):

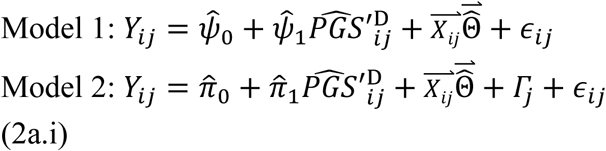

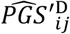: Normalized PGS constructed from the observed linear relationship between genotype and outcome

*Y*_*ij*_: Outcome for individual *i* in family *j*

*Γ*_*j*_: Family *j* fixed effect

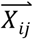: Vector of individual covariates 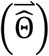 comprised of sex, age, and the first 10 principal components of genotype

Model 1 treats individuals as though they are unrelated whereas Model 2 compares siblings using a family fixed effect. In effect, Model 2 asks whether sibling differences in PGS translate into sibling differences in the outcome. Thus, Model 1 leverages covariation in *Y*_*ij*_ and 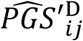 between individuals from different families while Model 2 leverages only covariation in *Y*_*ij*_ and 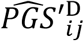 between pairs of individuals in the same family.

### 2b. Unresolved Questions

Table 1 displays results from Model 1 and Model 2 using data from the National Longitudinal Study of Adolescent to Adult Health (Harris 2013) for six phenotypes: educational attainment (Lee et al. 2018), cognitive ability (Lee et al. 2018), depressive symptoms (Turley et al. 2018), birth weight (Warrington et al. 2019), body mass index (Locke et al. 2015), and height (Wood et al. 2014). We further discuss the Add Health data and PGS construction (including links to the GWAS summary statistics used) in Sections A1 and A2 of the appendix.

**Table 1.** The association between polygenic score and observed trait for six phenotypes, within-families and between-families.

| | $\hat{\psi}_1$ | $\hat{\pi}_1$ | $\frac{\hat{\pi}_1}{\hat{\psi}_1}$ | $p(\hat{\psi}_1 = \hat{\pi}_1)$ |
| --- | --- | --- | --- | --- |
| Years of Schooling | 0.81** | 0.35** | 0.44 | 0 |
| Cognitive Ability | 3.16** | 1.7** | 0.54 | 0.02 |
| CESD Depression Index | 0.13** | 0.06 | 0.42 | 0.25 |
| Birth Weight | 3.1** | 3.26* | 1.05 | 0.91 |
| Body Mass Index | 2.01** | 2.34** | 1.17 | 0.59 |
| Height | 2.5** | 2.51** | 1 | 0.97 |

\* 0.05 \*\* 0.01 . All models control for sex, age, and the first 10 principal components of individual genotype. All model uses only individuals of European ancestry. Models without individual-fixed effects use a sample of unrelated individuals, whereas the family-fixed effect models use a sample of sibling pairs. The sample of unrelated individuals contains one randomly selected sibling from each sibling pair. All polygenic scores are both standardized within sample to be mean 0 and standard deviation 1. Cognitive ability is measured through the Peabody Picture Vocabulary Test during Wave 1 of Add Health, when respondents were approximately 16 years old. Birth weight is retrospectively reported by respondents' parents during Wave 1 of Add Health. Birth weight is retrospectively reported by respondents' parents during Wave 1 of Add Health. Years of schooling, CESD depression index, body mass index, and height are measured during Wave 4 of Add Health, when respondents were approximately 28 years old. Height is reported in centimeters, birth weight is reported in ounces, and cognitive ability is reported in IQ score points. The CESD depression index is normalized to be mean 0 and standard deviation 1. A traditional regression table is available in Section A12 of the appendix. P-values for the test that $\hat{\psi}_1 = \hat{\pi}_1$ were calculated through bootstrap resampling with replacement simulated with 1000 repetitions.

In Table 1, a one standard deviation increase in the educational attainment PGS is associated with over an additional .8 year of schooling between-families 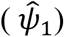 but less than half of that within-families 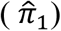. If we compare the six phenotypes, the relative size of 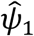 and 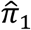 (captured by 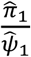) varies dramatically. For years of schooling and cognitive performance, bootstrapped p-values show that the differences seen within and between-family are statistically significant (i.e.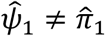). Moreover, these findings have been replicated by other researchers; a recent study using data from the United Kingdom found that PGS coefficients for cognitive traits were on average 60% greater between families than within-families (Selzam et al. 2019). They, like us, found no statistically significant differences between coefficients derived from within and between-family models for non-cognitive traits.

Why might this be the case? One possibility is that the between-family models are confounded while the within-family models capture the true causal effects of the PGS. Alternatively, it may be that some of the processes captured by GWAS function differently within-families versus between-families (for example, genetic nurturance). Answering this question requires a more formal treatment in order to better understand what underlying features may drive differences between 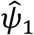 and 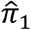 across phenotypes. Our theoretical model, which we develop below, suggests that, in addition to possible environmental confounding, bias in 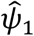 and 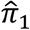 can depend on two novel parameters: (1) the underlying genetic correlation of direct and nurture effects and (2) the ratio of the SNP heritabilities for direct effects and nurture effects.

## 3. Theoretical Model

### 3a. Direct Genetic Effects and Genetic Nurture Effects

Historically, biosocial analyses have modeled complex traits as a function of both direct genetic effects and environment influences on an individual. Motivated by recent work highlighting the relevance of genetic nurture effects (Bates et al. 2018; Kong et al. 2018; Belsky et al. 2018; Wertz et al. 2018), we extend this model to include the genes of an individual’s parent. Thus, we assume outcome *Y*_*ij*_ is a function of individual *i*’s genotype, the genotypes of the parents in family *j*, and distinct individual-level and family-level environments. We choose to have a common effect of parental genetics at a given loci, instead of separate maternal and paternal effects, given the lack of strong empirical evidence of differences across parents (Kong et al. 2018).

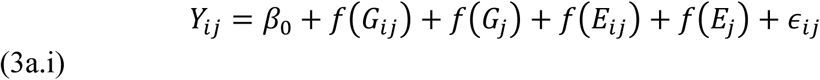

*f*(*G*_*ij*_): Effect of *i*′s genome on *Y*_*ij*_

*f*(*G*_*j*_): Effect of family *j*′s genome on *Y*_*ij*_

*f*(*E*_*ij*_): Effect of *i*′s environment on *Y*_*ij*_

*f*(*E*_*j*_): Effect of family *j*′s environment on *Y*_*ij*_

Note that here the environmental components *E*_*ij*_ and *E*_*j*_ are defined as the strictly non-genetic sources of variation in *Y*_*i*_. In other words, an environment influences a child’s *Y*_*i*_ irrespective of their or their parents’ genetic composition. Environmental features of a child’s environment, such as their family’s socioeconomic status, are captured in *f*(*G*_*j*_) if those environmental features were caused by their parents’ genes. We make three important assumptions to simplify the exposition of this model: no gene-environment interaction, no gene-environment correlation, and no assortative mating (see Section A3 of the appendix for a mathematical expression of these assumptions). We discuss the likely implications of a violation of these assumptions for our results at the end of Section 5a. While some of these assumptions are unlikely to hold true in the real world, we emphasize that goal of our theoretical model is not to create a model that perfectly describes reality but rather to formalize the ways in which social genetic effects influence GWASs, PGSs, and inevitably the interpretation of findings in the field of social science genomics. Our model, while simple, illustrates the key empirical phenomenon of interest; these higher-order features of the real world should not change the key implications derived from our model.

### 3b. True Polygenic Scores

Complex, behavioral traits are associated with many genes across the genome that simultaneously produce very small effects (Chabris et al. 2015; Visscher et al. 2017). To increase statistical power and simplify computation, researchers often summarize the relevant genetics of individual *i* into a single linear predictor called a PGS (Dudbridge 2013). This has become a widely utilized technique (Duncan et al. 2018) and relies on the assumptions that genetic effects are linear and additive. Recent meta-analyses of twin studies support the linear, additive model for genetic effects (Polderman et al. 2015). For the remainder of the paper, we approximate *f*(*G*_*ij*_) and *f*(*G*_*j*_) in our theoretical model (3a.i) using PGSs.

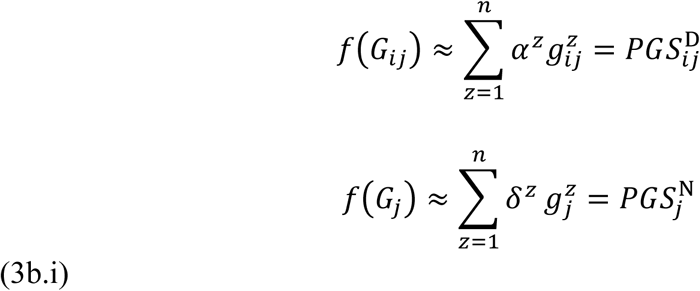

*α*^*z*^: True causal effect of a one allele change at *i*′s genetic loci *z* on *Y*_*ij*_

*δ*^*z*^: True causal effect of a one allele change at either parent in family *j*′s genetic loci *z* on *Y*_*ij*_

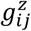: Total number of risk alleles at *i*′s genetic loci *z* (0, 1, or 2)

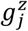: Total number of risk alleles at the parents in family *j*′s genetic loci *z* (0, 1, 2, 3 or 4)

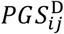: PGS constructed from the true causal linear effect of *i*’s genes on *Y*_*ij*_

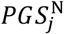: PGS constructed from the true causal linear effect of the parents in family *j*′s genes on *i*’s child′s *Y*_*ij*_

Notice that 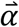 is the vector of causal allelic weights used to construct the true, underlying PGS for direct genetic effects. In the same vein, 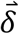 is the vector of causal allelic weights used to construct the true, underlying PGS for genetic nurture effects. Note that both 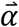 and 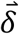 are structural parameters that are never empirically observed.

In Equation 3b.ii, we rewrite our theoretical model using PGSs.

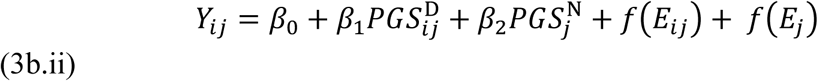

Notice that, because 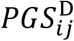 and 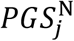 are not standardized, an individual’s value for 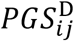 and 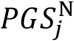 represents the true effect of their genes and their parents’ genes, respectively, on *Y*_*ij*_ *in the units of Y*_*ij*_. Thus, *β*_1_ and *β*_2_ are both equal to 1 by construction and Equation 3b.ii can equivalently be written without the *β*_1_ and *β*_2_ terms.

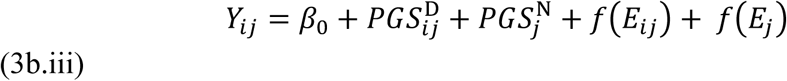

### 3c. Transmitted Genetic Nurture Alleles

The presence of social genetic effects, such as genetic nurture effects, will only bias GWAS estimates of direct genetic effects when a social or biological process induces a correlation between the genetics of an individual and the genetics of his or her relevant social relationships. In the case of genetic nurture effects, biological recombination acts as such a process; children randomly inherit a portion of each parent’s genome, leading to a mechanical correlation between parental genetics and child genetics. To capture the portion of the genetic nurture PGS that was transmitted to individual *i* in family *j* from their parents, we introduce a third PGS parameter that is absent from our formal model, 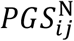.

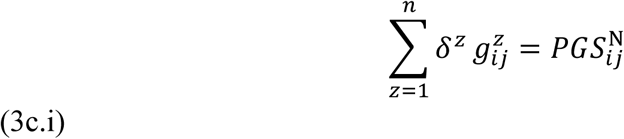

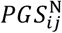: PGS constructed from the true causal linear effect of *i*’s genes on *i*’s child′s *Y*

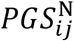 is constructed using aspects of both 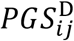 and 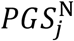, the two PGSs corresponding to the two causal sources of genetic effects present in our theoretical model. Like 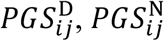 is constructed using *g*_*ij*_ (as opposed to *g*_*j*_) and therefore varies within-families. However, like 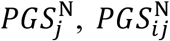 is constructed using the allelic weights 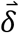, which correspond to genetic nurture effects (as opposed to direct genetic effects).

Notice that the relationship 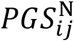 and 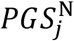 hinges on the relationship between *g*_*ij*_ and *g*_*j*_. Because the alleles transmitted from parent (*g*_*j*_) to child (*g*_*ij*_) are determined stochastically through genetic recombination, the expected value of the correlation between *g*_*ij*_ and *g*_*j*_ is known. In Section A4 of the appendix, we derive the expected value of the association between 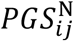 and 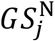.

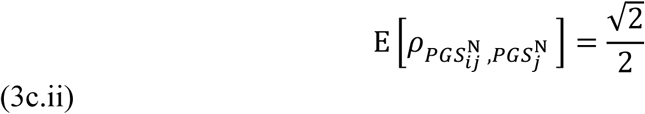

Note that this quantity does not vary between traits; this is due to the fact that same vector of allelic weights is used to construct both 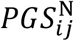 and 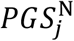 and differences are due entirely to the trait-independent recombination of parental alleles.

### 3d. The Relationship Between Direct Genetic Effects and Genetic Nurture Effects

While 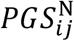 is absent from our underlying theoretical model, it provides the link between 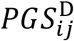 and 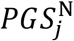 that allows for the presence of genetic nurture effects to distort GWAS results and subsequent PGS analysis. Thus, the most important components of our theoretical model are relationships between 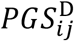 and 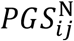 and between 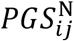 and 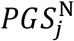. As we have seen above, 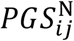 and 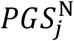 have a mechanical correlation that does not vary between traits as it depends only on genotype (and not allelic weights). However, any differences between 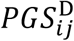 and 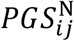 are a result of differences between allelic weights used to construct each PGS (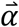 and 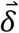, respectively). Because 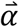 and 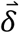 vary across traits, the extent to which 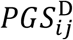 and 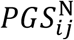 are correlated also varies between traits. We term the correlation between 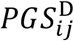 and 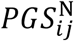 for a given trait the *direct-nurture genetic correlation*.

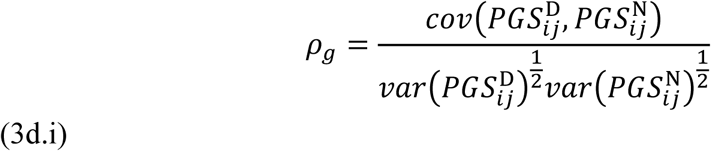

*ρ*_*g*_: Correlation between genetic nurture effects and direct genetic effects

In general, we expect the direct-nurture genetic correlation to be positive. Consider, for example, the case of educational attainment. Presumably some genetic pathways that lead a parent to create an environment conducive to their child succeeding in school will also have impacted the parent’s schooling. However, there may also exist some genetic pathways that contribute to a parent’s ability to create a positive educational environment for their children that do not influence the amount of educational attainment that the parent receives themselves. In the aggregate, we thus expect direct genetic effects and genetic nurture effects to have a correlation of less than 1. This value could in fact be negative, but, given the existing evidence suggests that *ρ*_*g*_ is typically positive (Kong et al. 2018; Bates et al. 2018; Belsky et al. 2018; Wertz et al. 2018), we focus on the case where the direct-nurture genetic correlation is bounded by 0 and 1.

### 3e. Underlying Allelic Weights

We now turn to a discussion of *α*^*z*^ and *δ* ^*z*^. In the course of this discussion, we will show that *ρ*_*g*_, a trait’s direct-nurture genetic correlation, is a critical structural parameter of our theoretical model. We first discuss *α*^*z*^. Allelic weights *α*^*n*^ (*n* ∈ {1 … *N*}) are taken from a distribution with variance *σ*. Without loss of generality, we define “risk” allele at each genetic loci *z* such that this distribution has mean 0. Turning to *δ*^*z*^, we assume that 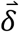 are drawn from an identical distribution as 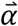, except with variance 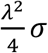. Note that this is related to the variance of the 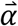 distribution but scaled by a parameter *λ*; this allows for the average effects sizes to differ between direct genetic effects and genetic nurture effects. Finally, we assume that genetic nurture effects and genetic nature effects have a similar genetic architecture such that 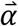 are 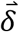 distributed in a similar way across genetic loci with respect to the allele frequency mean and variance. See Section A6 of the appendix for a formal treatment of this assumption.

The assumptions of our theoretical model entail that *λ* represents the ratio of the SNP heritabilities of genetic nurture effect and direct genetic effects (see Section A7 of the appendix). We call this value the *direct-nurture heritability ratio*.

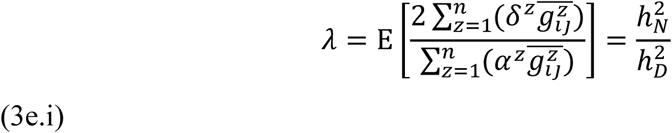

*λ*: Ratio of the SNP heritabilities of genetic nurture effects and direct genetic effects

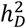: SNP heritability for direct genetic effects of *Y*_*ij*_

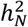: SNP heritability for genetic nurture effects of *Y*_*ij*_

Without loss of generality, we normalize all variables such that the variance of 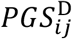 is equal to one.

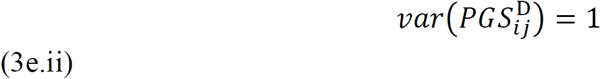

We can now use our theoretical model to derive the expectation of the variance of the remaining PGSs as a function of the direct-nurture heritability ratio (see Sections A8 and A9 of the appendix).

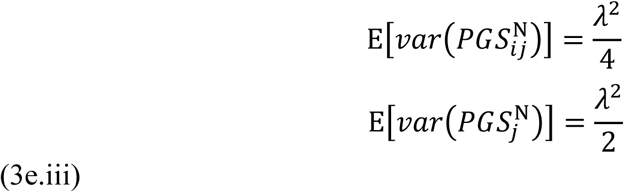

Now that we have fully identified our theoretical model, we can use it to gain insights into the implications of genetic nurture effects for GWAS and PGSs.

## 4. Analytic Results

### 4a. Observed PGS

Up until this point, all our work has been theoretical; we have defined the functional form of a set of causal relationships between underlying parameters of interest which are difficult to observe directly. We now transport our theoretical model into the messy real world and consider its implications for GWAS and subsequent PGSs. In reality, we observe not 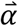 but 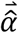. We then use this observed 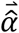 to construct not 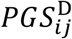 but 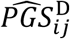, which, as we will see, contains information from both 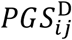 and 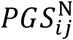.

We obtain 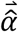 by fitting the following regression for *n* SNPs via GWAS.

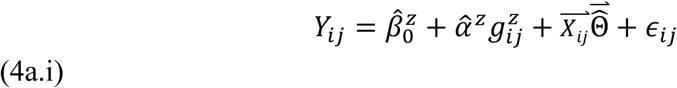

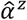: Allelic weight from the observed linear relationship of a one allele change at *i*′s *z*^*th*^ gene and *Y*_*ij*_

We can now plug in from our theoretical model (3b.iii).

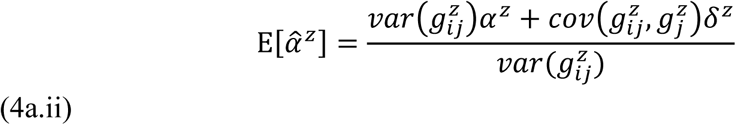

To further analyze this expectation, we can separate 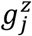 into the sum of alleles that were transmitted to *i* and the alleles that were not transmitted.

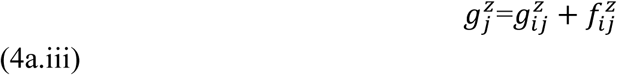

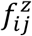: Total number of risk alleles at the parents in family *j*′s genetic loci *z* that were **not** transmitted to *i* (0, 1, or 2)

Thus yielding:

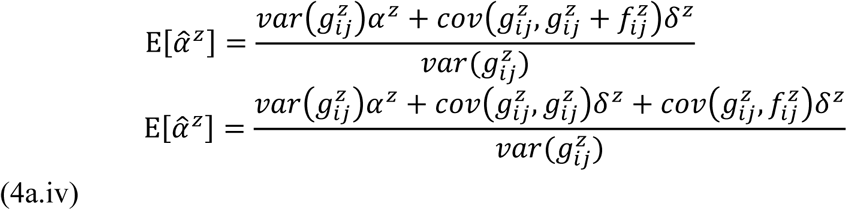

In the absence of assortative mating, transmitted alleles are uncorrelated with non-transmitted allele, meaning that 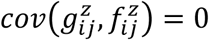.

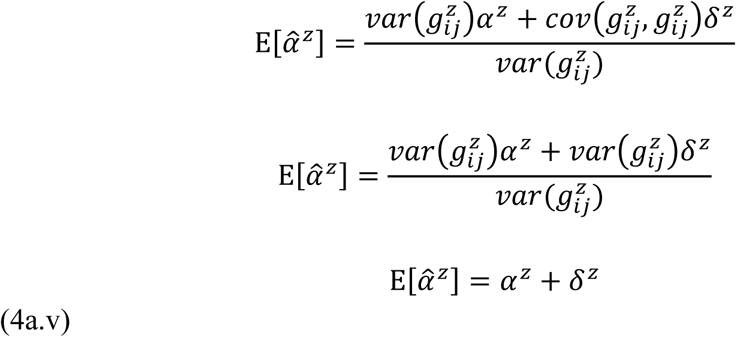

We can then use our observed vector of allelic weights for genetic nurture, 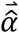, to construct our observed genetic nurture 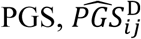.

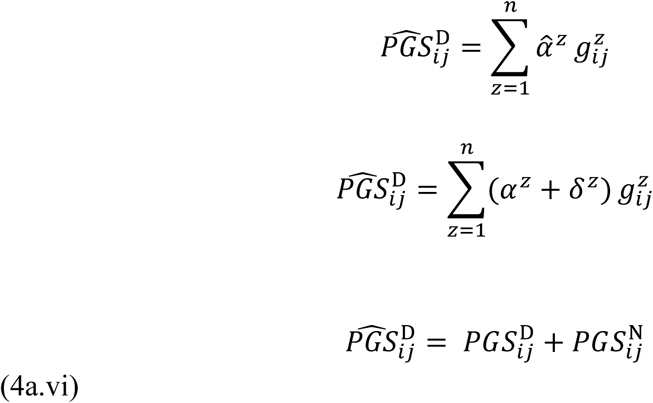

Finally, constructed PGSs are typically normalized within sample, so we convert 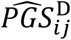 to 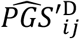.

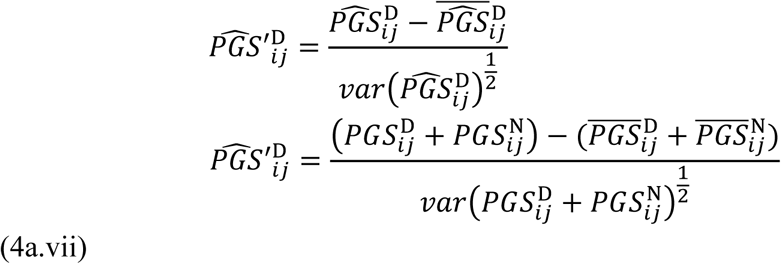

Thus, we have shown that the PGS derived from GWAS has information from both an individual’s PGS for direct genetic effects and genetic nurture effects. In particular, note that in (4a.v) the estimated weights intermingle both *α* and *δ*. As a result, the quantity typically used for analysis, 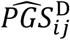, contains information about both the direct genetic effect and the genetic nurture effect (4a.vi).

### 4b. Between-Family Analyses

Our theoretical model has previously demonstrated that PGSs constructed from GWAS capture both the direct genetic effects and the genetic nurture effects of a given allele. We now explore the implications of this result for analyses using such PGSs, beginning with the between-family analysis (Model 1). We will see that the inclusion of genetic nurture effects biases direct genetic effect coefficients (away from zero).

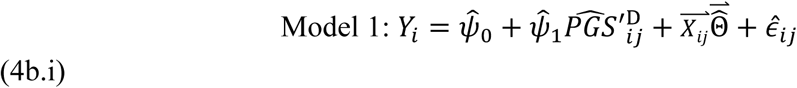

Now that we have derived 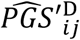, we can calculate the expected bias in 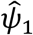. Recall that, because 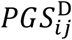 is unstandardized in our theoretical model (3b.iii), *Ψ*_1_and *π*_1_ are mechanically both equal to one. Thus, the expected value of 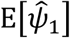 and 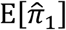 represents the expected inflation or deflation of estimated PGS coefficients (i.e. bias) in between-family and within-family analyses, respectively.

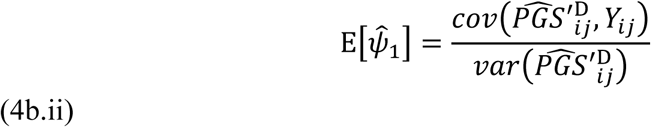

Note that 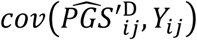 is given by true causal relationships from our theoretical model (3b.iii).

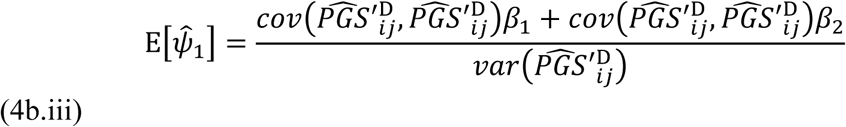

*β*_1_ and *β*_2_ from our theoretical model are equal to 1 by construction and therefore fall away.

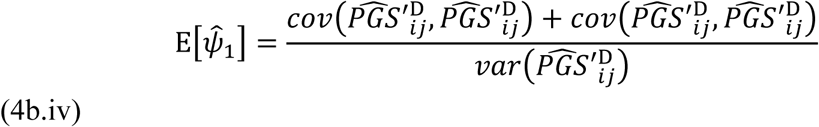

We solve (see Section A9 of the appendix for details) to obtain the magnitude of the bias in 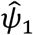.

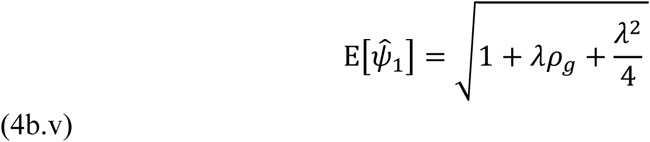

So we have shown that our observed estimates for 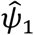 will be biased upwards by a factor 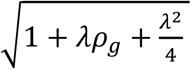. We will unpack this quantity further in the discussion but note that it depends on unobserved parameters.

### 4c. Within-Family Analyses

Let’s now turn to our within-family analysis (Model 2). We will see that the inclusion of genetic nurture effects biases direct genetic effect coefficients towards zero.

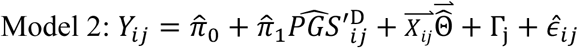

Or, equivalently:

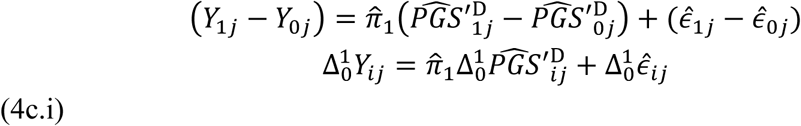

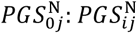 of sibling 0 in family *j*

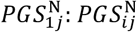 of sibling 1 in family *j*

As before, we begin by deriving the expected value of 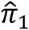.

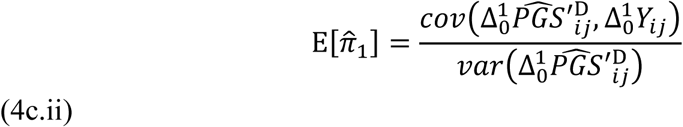

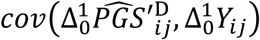 is given by true causal relationships from our theoretical model.

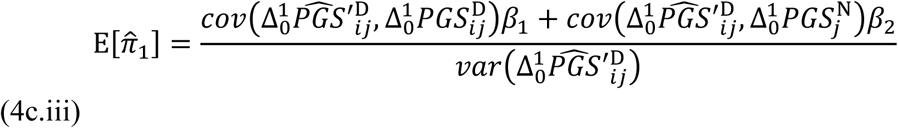

Again, *β*_1_ and *β*_2_ from our theoretical model are equal to 1 by construction and therefor fall away.

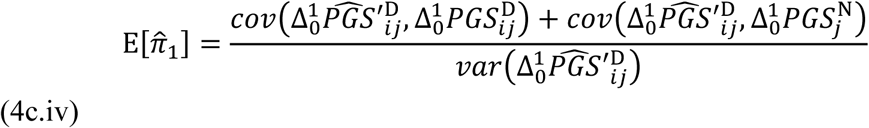

Notice that between siblings there is no variation in family genetic nurturing environment, meaning that 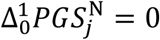.

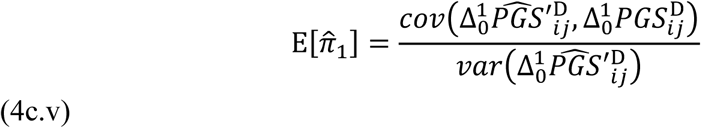

Now we just solve (see Section A11 of the appendix for details) to obtain the magnitude of the bias in 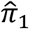.

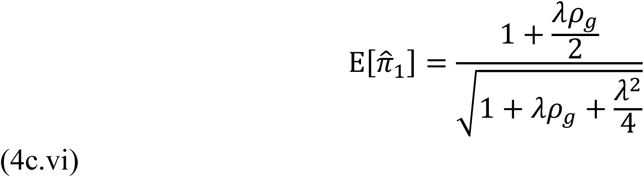

Recall that, because 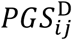 is unstandardized in our theoretical model (3b.iii), *π*_1_ is mechanically equal to one. Thus, the expected value of 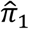 represents inflation or deflation of PGS coefficient estimates and is itself interpretable as measure of bias. Thus, we have shown that our observed estimates for 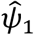 will be biased downwards by a factor of 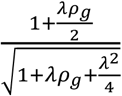. We discuss this quantity further in the discussion.

## 5. Discussion

### 5a. Bias

Our theoretical model illustrates that, in the presence of genetic nurture (i.e 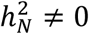), regression analyses using PGSs to estimate the effects of an individual’s genetics on their outcomes will suffer from bias. Between-family OLS models will be biased upwards by a factor of 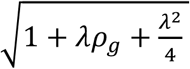 while within-family fixed effect models will be biased downwards by a factor of 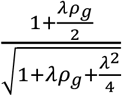. Figure 1 plots the bias in both OLS and family fixed effect regressions using PGSs as a function of various direct-nurture genetic correlations and direct-nurture heritability ratios.

**Figure 1.**
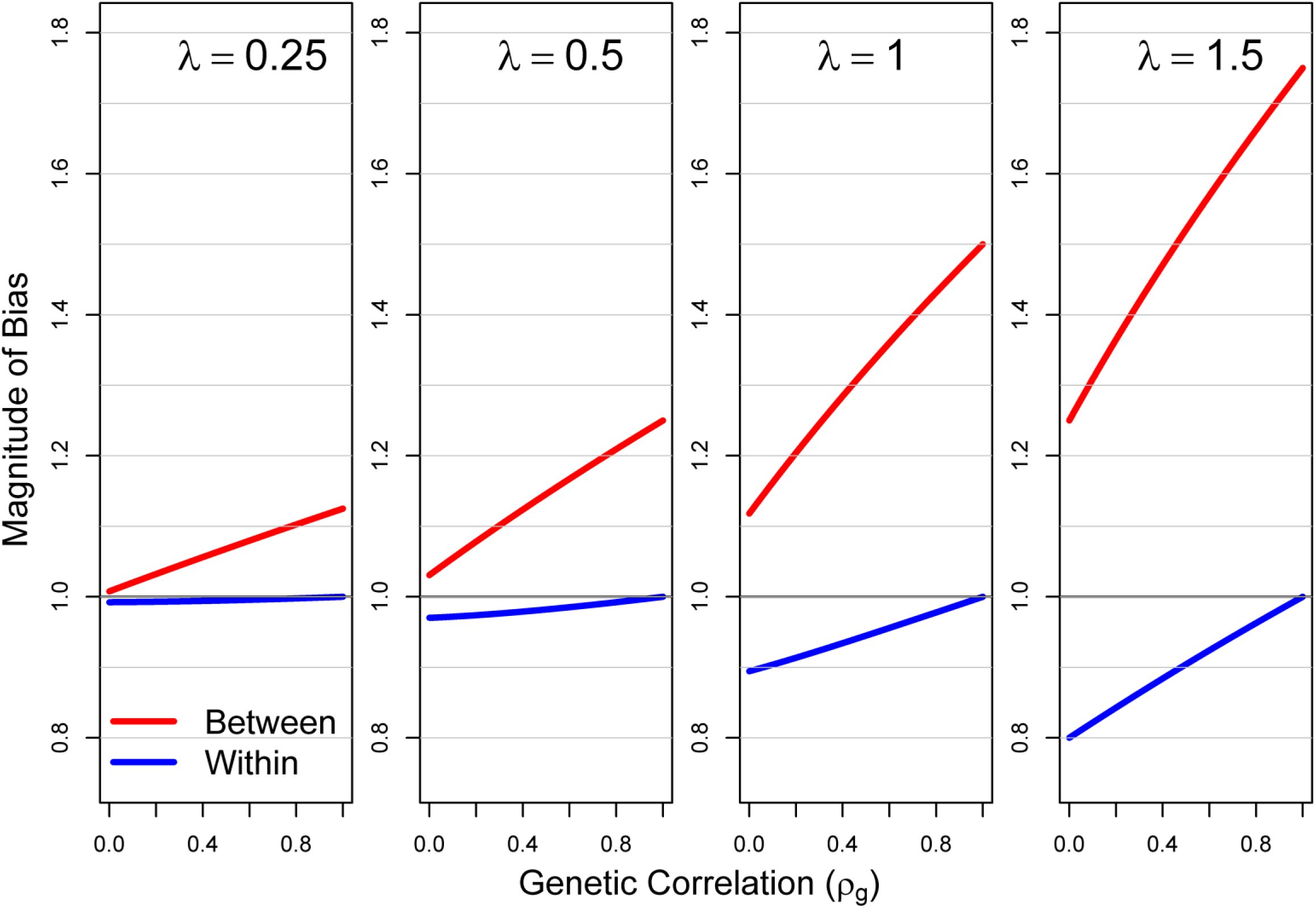
Bias Due to Genetic Nurture in Within and Between-family Regressions using Polygenic Scores. Grey line at y=1 represents no bias. *ρ*_*g*_ is the direct-nurture genetic correlation and *λ* is the direct-nurture heritability ratio. Results are derived analytically from a theoretical model.

The absence of bias is represented in Figure 1 by the horizontal line at y=1. For any given trait, bias is always larger in the between-family models. The magnitude of bias is a function of two parameters: *ρ*_*g*_, the trait’s direct-nurture genetic correlation, and *λ*, the trait’s direct-nurture heritability ratio. As *λ* increases, so does the magnitude of the bias. However, *ρ*_*g*_ has opposing effects on the bias within and between-families; as *ρ*_*g*_ increases, we note more downward within-family bias but more upward between-family bias. Thus, taken together, 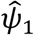 and 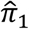 can provide a useful set of upper and lower bounds of the true causal effects.

There is an intuitive interpretation to the trends presented in Figure 1. In the between-family estimates, the inclusion of genetic nurture effects in 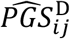 leads to an upward bias as it is capturing both differences in genetic composition between individuals *and* differences in the family environments between individuals that result from differences in their parents’ genetic composition. The larger that *ρ*_*g*_ is, the greater extent to which an individual with a beneficial allele for educational attainment reaps the reward from the same genes twice; first when their parents provide a more nurturing environment, and second when they themselves inherit the beneficial allele.

On the other hand, in family fixed effects models (within-family), the inclusion of genetic nurture in the observed PGS leads to downward bias when *ρ*_*g*_ is less than unity. This is because there are no differences in genetic nurture systematically driving differences in educational outcomes between siblings. Regardless of their own 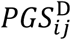, siblings have identical 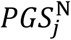 because they are in the same family *j*. Thus, any extent that genetic nurture effects causes 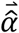 to diverge from 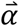 amounts to measurement error in the allelic weights and causes downward attenuation bias.

The insights from our theoretical model offer a partial explanation of the large differences in 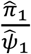 observed across the various phenotypes considered in Table 1. For traits like years of schooling, cognitive ability, and (more speculatively) depression, where indirect genetic effects likely play an important role (i.e. large *λ*), we see large differences between 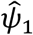 and 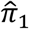 (i.e 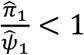). On the other side of the coin, for traits like body mass index and height, where most of the genetic contribution is likely to be direct (i.e. small *λ*), we see almost no difference between 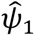 and 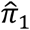 (i.e 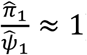). Turning back to Figure 1, you can see this coincides with what our theoretical model would have predicted. Further, our model suggests that the differences in 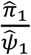observed between years of schooling, cognitive ability, and depression may be a function of differences in *ρ*_*g*_ between the traits.

Recall that these theoretical results are based on several simplifying assumptions: no gene-environment interaction, no gene-environment correlation, and no assortative mating. These three assumptions are unlikely to hold for most complex traits studied of interest to the social and biological sciences. Nonetheless, we can use the results from this simplified model to begin to probe how such violations might influence our results. Consider the case of positive genetic assortative mating, which exists for many of the traits considered in this paper (Yengo, Robinson, et al. 2018). A correlation between maternal and paternal genetics induces a positive correlation between transmitted and non-transmitted parental alleles shown in (4a.iv). This is because non-transmitted maternal alleles would be correlated with transmitted paternal alleles and vice versa. Thus, 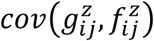 is no longer equal to zero and does not fall out of our equation. In such a case, the expected value of a GWAS allelic weight, shown in (4a.v), would include information about both non-transmitted genetic nurture effects in addition to the direct genetic effects and the transmitted genetic nurture effects. Thus, the bias documented in PGS analyses will increase as positive genetic assortative mating increases.

Next, we consider the case of gene-environment correlation. It is important to note that the bias that genetic nurture may cause in GWAS and PGS results from a special case of gene-environment correlation (i.e. a gene-environment correlation that mechanically exists due to the correlation of genetics between parents and their children induced though genetic inheritance). Thus, in our theoretical model’s specification, gene-environment correlation only exists when environmental features relevant to an outcome are correlated with an individual’s genetics after accounting for their parents’ genetics. When such a gene-environment correlation exists, GWAS results and PGSs become biased in exactly the same way as they do from genetic nurture effects alone. For example, the existence of positive gene-environment correlation effectively increases *λ* (as additional outcome variance explained by a non-genetic nurture environmental component), therein increasing the magnitude of the bias.

Finally, the case of gene-environment interaction is difficult to consider more generally, as it would vary as a function of the magnitude, direction, and the pathways of the interaction. Thus, the effects of gene-environment interactions on how genetic nurture effects influence GWAS and PGSs remain uncertain.

### 5b. Direct-Nurture Genetic Correlation and Heritability Ratio

Numerous GWAS have been conducted in the last decade (Visscher et al. 2017; Mills and Rahal 2019). Nonetheless, to our knowledge, no GWAS has been conducted in human populations that independently identifies the direct genetic effects and the genetic nurture effects for a complex trait (a notable exception is GWAS on maternal influences on child birthweight (Beaumont et al. 2018; Warrington et al. 2019), though this indirect genetic affect does not seem to be social in nature). Thus, for virtually all complex traits, little is known about the parameters of interest identified in our models: the direct-nurture genetic correlation and the direct-nurture heritability ratio (*ρ*_*g*_ and *λ* respectively). Critically, the two parameters are readily estimable with existing data and methods (which we discuss further in 5d).

A better understanding of *ρ*_*g*_ and *λ* would offer value beyond aiding in the comparison of results from within-family and between-family regressions. The genetic pathways discovered in GWASs and summarized in PGSs offer researchers a puzzle to unpack (Freese 2018). Understanding why some phenotypes have strong versus weak social genetic effects, or why the link between direct genetic and social genetic influence are more versus less linked, could help researchers glean insight into the underlying mechanisms at play.

Moreover, it would be interesting to understand how *ρ*_*g*_ and *λ* are influenced by the social environment. For example, while gene-environment interaction studies have been the conventional way to understand how the environment moderates the influence of genetics, recent work has proposed a genetic correlation-environment interaction study (Wedow et al. 2018). In a genetic correlation-environment interaction study framework, the social environment can transform the genetic link between two traits. Exploring how the environment shapes *ρ*_*g*_would be a special case of a genetic correlation-environment interaction study where the two traits influence the same phenotype (directly and socially). Social policymakers might prefer a low *ρ*_*g*_ for valued life outcomes like educational attainment to reduce the accumulation of inequality across generations.

While the specific social, physical, or economic factors that moderate *ρ*_*g*_ and *λ* for various traits remains to be explored empirically, there may exist *a priori* reasons to suspect certain environmental modifiers. For example, let us say that individual variation in height is a function of both direct genetic effects that shape physiological development and genetic nurture effects that influence access to socioeconomic and nutritional resources during childhood (thereby reducing the likelihood of stunting). If there is a large casual effect of height on socioeconomic status, we’d expect individuals with a greater genetic predisposition for height (i.e. a high 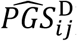)to be more likely to attain a higher socioeconomic position where their children to have access to nutritional and health resources (i.e. a high 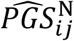) resulting in a positive *ρ*_*g*_. However, this relationship could be modified by environmental features; if, for instance, the causal effect of height on social status is due to labor force discrimination, outlawing the use of height for employment decisions would uncouple 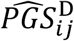 and 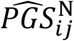 and reduce *ρ*_*g*_. Alternatively, a social policy that provides adequate healthcare and nutrition to all children could effectively eliminate stunting and undo the relationship between parental genetics for height altogether, forcing both *ρ*_*g*_ and *λ* to zero.

### 5c. Implications for the Use of Polygenic Scores

While within-family analyses have demonstrated that many PGSs do have significant casual signal for direct genetic effects, comparing the results from within-family analyses to results from between-family analyses is complicated by the presence of genetic nurture effects. To what extent do existing PGSs capture direct genetic effects, genetic nurture effects, and socioeconomic or geographic confounding? Until we better understand *ρ*_*g*_ and *λ* for a wide variety of traits, our ability to use within-family analyses to validate between-family discoveries will be limited. Analyses using PGSs should be interpreted accordingly.

Even in the absence of confounding due to population stratification, the observation that existing PGSs likely have genetic nurture components complicates their use and interpretation. Say, for instance, a researcher wonders whether there exists moderation of the association between an individual’s PGS and their educational attainment as a function of school-level socioeconomic status (Trejo et al. 2018). Because a component of the PGS is capturing the benefit of having a parent with higher educational attainment and (in turn a higher socioeconomic status), any detected gene-environment interaction lacks a clear interpretation. It might be the case that household socioeconomic status interacts with school-level socioeconomic status in shaping educational attainment, or alternatively it could be that an individual’s genetic composition interacts with school-level socioeconomic status in shaping educational attainment. These two different results have very different theoretical and practical implications but are indistinguishable in analyses using existing PGSs, which contain both direct genetic effects and genetic nurture effects.

### 5d. Future Research

The models constructed in this paper highlight key areas for future research in the field of social science genomics. Across a range of complex phenotypes, there is much work to be done towards separating out the genetics nurture effects from direct genetic effects. Utilizing random cage-mate assignment in mice, a recent study in mice conducted a social genetic effects GWAS and direct genetic effects GWAS in parallel and identified statistically significant genome wide social genetic effect loci for 16 phenotypes (Baud et al. 2018). For these 16 phenotypes, the mean social-direct genetic correlation was 0.53 and the mean social-direct heritability ratio was 1.29. Crucially, social genetic effects arise from partially different loci as direct genetic effects and can have effects of differing magnitudes or directions at the same loci.

Unfortunately, social relationships are often not randomly assigned in human populations. Nonetheless, existing GWAS methods could be modified to use dyads of parents and their children. For example, a social genetic effects GWAS could be conducted by controlling for child’s genetics in a GWAS of parental genetics on child phenotype. Alternatively, a GWAS conducted using variation only amongst sibling pairs would provide information on direct genetic pathways untainted by confounding or genetic nurture effects. Results from such a sibling GWAS might then be used to back out information about the genetic nurture effects from existing GWAS results of unrelated individuals. Nonetheless, obtaining large samples of parent-child or sibling pairs might prove challenging for many complex phenotypes. If statistical power is a problem, methods such as LD score regression (Bulik-Sullivan et al. 2015) could be used with smaller samples to identify estimates of *ρ*_*g*_ and *λ*. Indeed, having estimates of *ρ*_*g*_ and *λ* for a trait would allow researchers to correct for bias in between-family and within-family analyses that use PGSs by dividing observed regression coefficients by the quantities displayed in (4b.v) and (4c.vi), respectively.

It is also possible that the direct effects and social genetic effects are not independent. Imagine, for example, that parents with a higher educational attainment PGSs tend to invest more heavily in their children with higher PGSs than do parents with lower PGSs (differential investment in low birth weight children has been observed by across socioeconomic lines (Hsin 2012; Restrepo 2016)). In such a case, the effects of parental genetics on a child would vary as a function as their child’s genetics. Research designs that investigate the existence of interaction between direct genetic effects and genetic nurture effects may prove a fruitful avenue for future inquiry.

Finally, work should be done to extend our framework to other social genetic effects within-families, such as between sibling pairs. We chose to start with genetic nurture effects because, unlike sibling effects, genetic nurture effects are likely to be unidirectional, with the causal effect pointing from parent to child, making them more straightforward to model (effects between siblings are likely reciprocal) (Kong et al. 2018) and because parental effects generalize to all families, not just those with multiple children. Nonetheless, social genetic effects between siblings may also complicate the interpretation of GWAS results and warrant attention.

### 5d. Conclusion

In summary, we formalize a theoretical model for additive direct genetic effects and genetic nurture effects and show that, unlike bias from other confounders, the presence of genetic nurture can bias coefficients from between-family and within-family regressions using PGSs. While within-family analyses that compare siblings using family fixed effects are considered the gold-standard, they are not without their own complications. Even if we were able to run a GWAS on an infinitely large sample using the methods in practice today, the presence of confounding social genetic effects would mean that it would be impossible to obtain precise estimates of the causal effect of an individual’s genes on their life outcomes. Until GWAS can be conducted controlling for parental genotype, models using PGSs may be biased by genetic nurture effects. Obtaining estimates of a trait’s direct-nurture genetic correlation and direct-nurture heritability ratio may allow researchers to correct for such a bias.

## Acknowledgements

We wish to thank Kathleen Mullan Harris and the Add Health staff for access to the restricted-use genetic data. We are grateful to Patrick Turley, Dalton Conley, attendees of the Stanford Genetics & Social Science journal club, seminar participants at the Integrating Genetics and Society 2018 Conference, and two anonymous referees for helpful comments. This work has been supported by the Russell Sage Foundation and the Ford Foundation under Grant No. 96-17-04, the National Science Foundation under Grant No. DGE-1656518, and by the Institute of Education Sciences under Grant No. R305B140009. Any opinions expressed are those of the authors alone and should not be construed as representing the opinions of any foundation.

