## Supplementary material for "Genetic Nature or Genetic Nurture? Quantifying Bias in Analyses Using Polygenic Scores": Online Appendix

#### A1. Add Health

The National Longitudinal Study of Adolescent to Adult Health (Add Health) is a nationally representative cohort drawn from a probability sample of 80 high schools and 52 middle schools in roughly 80 US communities, and representative of schools in the United States in 1994–95 with respect to region, urban setting, school size, school type, and race or ethnic background. About 15,000 Add Health respondents (or 96%) consented to genotyping during the Wave 4 interview in 2008-09 for purposes of approved Add Health Wave 4 research. Of those who consented to genotyping, ~12,000 (or 80%) agreed to have their DNA archived for future testing. DNA extraction and genotyping were conducted on this archive sample using two platforms (Illumina Omni1 for siblings and the Illumina Omni2.5 for unrelated individuals). Quality control procedures were performed on the genetic samples collected yielding genetic data from ~10,000 individuals on 609,130 overlapping SNPs.

We focus most analyses on a set of genetically homogenous respondents of European descent. We restrict our sample to only European ancestry individuals because differences in linkage disequilibrium and allele frequencies that exist across ancestral groups complicate the interpretation of PGS–phenotype associations (Martin et al. 2017). Although we recognize the importance of research in more diverse samples, Add Health does not contain large enough samples of any ancestral group other than European ancestry to conduct well-powered within family analyses. Fortunately, the theoretical results regarding how genetic nurturance can induce bias in GWASs and the PGSs constructed using their summary statistics, which we emphasize is the core contribution of our paper, applies to genetic analyses conducted in all ancestry groups (although the relevant underlying $\lambda$ and $\rho_g$ parameters may vary between such groups due to environmental differences). Add Health also contains a variety of data on students' academic performance, personal characteristics measured in adolescence (cognitive ability, personality characteristics, professional aspirations, physical health and functioning, etc.).

#### A2. European Ancestry Identification and Polygenic Score Construction

To identify a sample of European-ancestry respondents, we calculated the first two principal components participants with known ancestries from the 1000 Genomes Project. We then projected Add Health individuals onto those principal components, obtaining the loadings of each Add Health respondent on each PC. We use these loading to assign each individual to one of five superpopulations in the 1000 Genomes data: European, African, East Asian, South Asian and Admixed and restrict our sample to only individuals of European ancestry. In this sub-sample, calculated PCs that we use as controls in our polygenic score regression analyses.

Polygenic scores (PGSs) were created using SNPs in the Add Health genetic database that were matched to SNPs with reported results in a GWAS. We also pruned all SNPs where the risk allele identified via GWAS could not be readily identified in the Add Health genetic database. For each SNP, a loading was calculated as the number of trait-associated alleles multiplied by the effect size estimated in the original GWAS. SNPs with relatively large p-values will have small effects (and thus be down weighted in creating the composite), so we do not impose a p-value threshold. Loadings were summed across the SNP set to calculate the polygenic score. The scores are standardized within sample to have a mean of 0 and standard deviation of 1. PGS generated from GWAS that included Add Health were constructed with from summary statistics with Add Health removed. Links to the summary statistics used to construct each score are provided below:

Body mass index: https://portals.broadinstitute.org/collaboration/giant/images/c/c8/Meta-analysis_Locke_et_al%2BUKBiobank_2018_UPDATED.txt.gz
Height: https://portals.broadinstitute.org/collaboration/giant/images/6/63/Meta-analysis_Wood_et_al%2BUKBiobank_2018.txt.gz
Own birth weight:
http://mccarthy.well.ox.ac.uk/publications/2019/EggBirthWeight_NatureGenetics/Fetal_BW_European_meta.NG2019.txt.gz
Educational attainment*: http://ssgac.org/documents/MTAG_EA.to10K.txt
Cognitive performance*: http://ssgac.org/documents/MTAG_CP.to10K.txt
Depressive symptoms*: http://ssgac.org/documents/MTAG_DEP_CLUMPED.to10K.txt

*Note that the Social Science Genomics Association Consortium phenotypes (i.e. educational attainment, cognitive performance, and depressive symptoms) used 23&Me data in their published GWAS. While our polygenic scores were generated using the full set of summary statistics, the publicly available data (linked to above) contain only the 10K SNPs with the lowest p-values in compliance with 23&Me’s data sharing policies.

**A3. Simplifying Assumptions of Structural Model**

We make several simplifications to simplify the exposition of the structural model. In defining our structural model, we have already assumed away gene-environment interactions (notice the lack of an interaction term between genes and environment). We further assume no gene-environment correlation.

$$cov(PGS_{ij}^{\mathrm{D}}, E_{ij}) = cov(PGS_{j}^{\mathrm{N}}, E_{j}) = 0$$

Finally, we assume no assortative mating. This means that the polygenic scores of parents are uncorrelated.

$$PGS_j^{\mathrm{N}} = \sum_{z=1}^{n} \delta^z g_j^z = \sum_{z=1}^{n} \delta^z (g_j^{zm} + g_j^{zp}) = \sum_{z=1}^{n} \delta^z g_j^{zm} + \sum_{z=1}^{n} \delta^z g_j^{zp} = PGS_j^{\mathrm{Nm}} + PGS_j^{\mathrm{Np}}$$

$$var\left(PGS_j^{\mathrm{N}}\right) = var\left(PGS_j^{\mathrm{Nm}} + PGS_j^{\mathrm{Np}}\right) = var\left(PGS_j^{\mathrm{Nm}}\right) + var\left(PGS_j^{\mathrm{Np}}\right) = 2var\left(PGS_{ij}^{\mathrm{N}}\right)$$

$g_j^{zm}$: Total number of major alleles at the mother in family $j'$s genetic loci $z$ $(0, 1, \text{or } 2)$

$g_j^{zp}$: Total number of major alleles at the father in family $j'$s genetic loci $z$ $(0, 1, \text{or } 2)$

$PGS_j^{\mathrm{Dm}}$: PGS constructed from the true causal linear effect of the mother in family $j'$s genes on $Y_{ij}$

$PGS_j^{\mathrm{Dp}}$: PGS constructed from the true causal linear effect of the father in family $j'$s genes on $Y_{ij}$

This also entails that a child's PGS is related to their parent's and sibling's PGS only directly through the genes they receive through the process of recombination. Thus, the expected value of the correlation between the same PGSs for all parent-child and sibling-sibling pairs is .5.

$$\mathrm{E}\left[\rho_{PGS_{0j}^{\mathrm{D}}, PGS_{1j}^{\mathrm{D}}}\right] == \mathrm{E}\left[\rho_{PGS_j^{\mathrm{Dm}}, PGS_{0j}^{\mathrm{D}}}\right] = \mathrm{E}\left[\rho_{PGS_j^{\mathrm{Dm}}, PGS_{1j}^{\mathrm{D}}}\right] = \mathrm{E}\left[\rho_{PGS_j^{\mathrm{Np}}, PGS_{0j}^{\mathrm{D}}}\right] = \mathrm{E}\left[\rho_{PGS_j^{\mathrm{Np}}, PGS_{1j}^{\mathrm{D}}}\right] = .5$$

$$\mathrm{E}\left[\rho_{PGS_{0j}^{\mathrm{N}}, PGS_{1j}^{\mathrm{N}}}\right] = \mathrm{E}\left[\rho_{PGS_j^{\mathrm{Nm}}, PGS_{0j}^{\mathrm{N}}}\right] = \mathrm{E}\left[\rho_{PGS_j^{\mathrm{Nm}}, PGS_{1j}^{\mathrm{N}}}\right] = \mathrm{E}\left[\rho_{PGS_j^{\mathrm{Np}}, PGS_{0j}^{\mathrm{N}}}\right] = \mathrm{E}\left[\rho_{PGS_j^{\mathrm{Np}}, PGS_{1j}^{\mathrm{N}}}\right] = .5$$

**A4. Correlation between $PGS_{ij}^{\mathrm{N}}$ and $PGS_{j}^{\mathrm{N}}$**

$$\rho_{PGS_{ij}^{\mathrm{N}},PGS_{j}^{\mathrm{N}}} = \frac{cov\left(PGS_{ij}^{\mathrm{N}},\ PGS_{j}^{\mathrm{N}}\right)}{var\left(PGS_{ij}^{\mathrm{N}}\right)^{\frac{1}{2}} var\left(PGS_{j}^{\mathrm{N}}\right)^{\frac{1}{2}}}$$

$$\rho_{PGS_{ij}^{\mathrm{N}},PGS_{j}^{\mathrm{N}}} = \frac{cov\left(PGS_{ij}^{\mathrm{N}},\ PGS_{j}^{\mathrm{N_m}} +\ PGS_{j}^{\mathrm{N_p}}\right)}{var\left(PGS_{ij}^{\mathrm{N}}\right)^{\frac{1}{2}} var\left(PGS_{j}^{\mathrm{N_m}} +\ PGS_{j}^{\mathrm{N_p}}\right)^{\frac{1}{2}}}$$

$$\rho_{PGS_{ij}^{\mathrm{N}},PGS_{j}^{\mathrm{N}}} = \frac{cov\left(PGS_{ij}^{\mathrm{N}}, PGS_{j}^{\mathrm{N_m}}\right) + cov\left(PGS_{ij}^{\mathrm{N}}, PGS_{j}^{\mathrm{N_p}}\right)}{var\left(PGS_{ij}^{\mathrm{N}}\right)^{\frac{1}{2}} [var\left(PGS_{j}^{\mathrm{N_m}}\right) + var\left(PGS_{j}^{\mathrm{N_p}}\right) + 2cov(PGS_{j}^{\mathrm{N_m}},\ PGS_{j}^{\mathrm{N_p}})]^{\frac{1}{2}}}$$

$$\rho_{PGS_{ij}^{\mathrm{N}},PGS_{j}^{\mathrm{N}}} = \frac{\rho_{,PGS_{ij}^{\mathrm{D}},PGS_{j}^{\mathrm{D_m}}} var\left(PGS_{ij}^{\mathrm{N}}\right)^{\frac{1}{2}} \left(PGS_{j}^{\mathrm{N_m}}\right)^{\frac{1}{2}} + \rho_{,PGS_{ij}^{\mathrm{D}},PGS_{j}^{\mathrm{D_p}}} var\left(PGS_{ij}^{\mathrm{N}}\right)^{\frac{1}{2}} \left(PGS_{j}^{\mathrm{N_p}}\right)^{\frac{1}{2}}}{var\left(PGS_{ij}^{\mathrm{N}}\right)^{\frac{1}{2}} [var\left(PGS_{j}^{\mathrm{N_m}}\right) + var\left(PGS_{j}^{\mathrm{N_p}}\right) + 0]^{\frac{1}{2}}}$$

$$\mathrm{E}\left[\rho_{PGS_{ij}^{\mathrm{N}},PGS_{j}^{\mathrm{N}}}\right] = \mathrm{E}\left[\frac{\rho_{,PGS_{ij}^{\mathrm{D}},PGS_{j}^{\mathrm{D_m}}} var\left(PGS_{ij}^{\mathrm{N}}\right)^{\frac{1}{2}} \left(PGS_{j}^{\mathrm{N_m}}\right)^{\frac{1}{2}} + \rho_{,PGS_{ij}^{\mathrm{D}},PGS_{j}^{\mathrm{D_p}}} var\left(PGS_{ij}^{\mathrm{N}}\right)^{\frac{1}{2}} \left(PGS_{j}^{\mathrm{N_p}}\right)^{\frac{1}{2}}}{var\left(PGS_{ij}^{\mathrm{N}}\right)^{\frac{1}{2}} [var\left(PGS_{j}^{\mathrm{N_m}}\right) + var\left(PGS_{j}^{\mathrm{N_p}}\right) + 0]^{\frac{1}{2}}}\right]$$

$$\mathrm{E}\left[\rho_{PGS_{ij}^{\mathrm{N}},PGS_{j}^{\mathrm{N}}}\right] = \frac{.5\lambda^2 + .5\lambda^2}{\lambda[\lambda^2 + \lambda^2]^{\frac{1}{2}}}$$

$$\mathrm{E}\left[\rho_{PGS_{ij}^{\mathrm{N}},PGS_{j}^{\mathrm{N}}}\right] = \frac{\sqrt{2}}{2}$$

### A5. Correlation between $\boldsymbol{PGS_{ij}^{\mathrm{D}}}$ and $\boldsymbol{PGS_{j}^{\mathrm{N}}}$

$$\rho_{PGS_{ij}^{\mathrm{D}},PGS_{j}^{\mathrm{N}}} = \frac{cov\left(PGS_{ij}^{\mathrm{D}},\ PGS_{j}^{\mathrm{N}}\right)}{var\left(PGS_{ij}^{\mathrm{D}}\right)^{\frac{1}{2}} var\left(PGS_{j}^{\mathrm{N}}\right)^{\frac{1}{2}}}$$

$$\rho_{PGS_{ij}^{\mathrm{D}},PGS_{j}^{\mathrm{N}}} = \frac{cov\left(PGS_{ij}^{\mathrm{D}},\ PGS_{j}^{\mathrm{N_m}} +\ PGS_{j}^{\mathrm{N_p}}\right)}{var\left(PGS_{ij}^{\mathrm{D}}\right)^{\frac{1}{2}} var\left(PGS_{j}^{\mathrm{N_m}} +\ PGS_{j}^{\mathrm{N_p}}\right)^{\frac{1}{2}}}$$

$$\rho_{PGS_{ij}^{\mathrm{D}},PGS_{j}^{\mathrm{N}}} = \frac{cov\left(PGS_{ij}^{\mathrm{D}}, PGS_{j}^{\mathrm{N_m}}\right) + cov\left(PGS_{ij}^{\mathrm{D}}, PGS_{j}^{\mathrm{N_m}}\right)}{var\left(PGS_{ij}^{\mathrm{D}}\right)^{\frac{1}{2}} [var\left(PGS_{j}^{\mathrm{N_m}}\right) + var\left(PGS_{j}^{\mathrm{N_p}}\right) + 2cov(PGS_{j}^{\mathrm{N_m}},\ PGS_{j}^{\mathrm{N_p}})]^{\frac{1}{2}}}$$

$$\rho_{PGS_{ij}^{\mathrm{D}},PGS_{j}^{\mathrm{N}}} = \frac{\rho_{,PGS_{ij}^{\mathrm{D}},PGS_{j}^{\mathrm{N_m}}} var\left(PGS_{ij}^{\mathrm{D}}\right)^{\frac{1}{2}} \left(PGS_{j}^{\mathrm{N_m}}\right)^{\frac{1}{2}} + \rho_{,PGS_{ij}^{\mathrm{D}},PGS_{j}^{\mathrm{N_m}}} var\left(PGS_{ij}^{\mathrm{N}}\right)^{\frac{1}{2}} \left(PGS_{j}^{\mathrm{N_p}}\right)^{\frac{1}{2}}}{var\left(PGS_{ij}^{\mathrm{D}}\right)^{\frac{1}{2}} [var\left(PGS_{j}^{\mathrm{N_m}}\right) + var\left(PGS_{j}^{\mathrm{N_p}}\right) + 0)]^{\frac{1}{2}}}$$

$$\mathrm{E}\left[\rho_{PGS_{ij}^{\mathrm{D}},PGS_{j}^{\mathrm{N}}}\right] = \mathrm{E}\left[\frac{\rho_{,PGS_{ij}^{\mathrm{D}},PGS_{j}^{\mathrm{N_m}}} var\left(PGS_{ij}^{\mathrm{D}}\right)^{\frac{1}{2}} \left(PGS_{j}^{\mathrm{N_m}}\right)^{\frac{1}{2}} + \rho_{,PGS_{ij}^{\mathrm{D}},PGS_{j}^{\mathrm{N_m}}} var\left(PGS_{ij}^{\mathrm{N}}\right)^{\frac{1}{2}} \left(PGS_{j}^{\mathrm{N_p}}\right)^{\frac{1}{2}}}{var\left(PGS_{ij}^{\mathrm{D}}\right)^{\frac{1}{2}} [var\left(PGS_{j}^{\mathrm{N_m}}\right) + var\left(PGS_{j}^{\mathrm{N_p}}\right) + 0)]^{\frac{1}{2}}}\right]$$

$$\mathrm{E}\left[\rho_{PGS_{ij}^{\mathrm{D}},PGS_{j}^{\mathrm{N}}}\right] = \frac{.5\rho\lambda + .5\rho\lambda}{[\lambda^2 + \lambda^2]^{\frac{1}{2}}}$$

$$\mathrm{E}\left[\rho_{PGS_{ij}^{\mathrm{D}},PGS_{j}^{\mathrm{N}}}\right] = \frac{\sqrt{2}}{2}\ \rho$$

### A6. Underlying Allelic Weights

Specifically, we assume:

$$\rho_{\vec{\alpha},\overrightarrow{\bar{g}_{ij}^{z}}} = \rho_{\vec{\delta},\overrightarrow{\bar{g}_{ij}^{z}}}$$

$$\rho_{\vec{\alpha}^2,\overrightarrow{var}(g_{ij}^{z})} = \rho_{\vec{\delta}^2,\overrightarrow{var}(g_{ij}^{z})}$$

$\bar{g}_{ij}^{z}$: Population mean risk allele frequency count at genetic loci $z$

$\overrightarrow{var}\left(g_{ij}^{z}\right)$ : Population variance of risk allele count frequency at genetic loci $z$

$\rho_{\vec{\alpha},\overrightarrow{\bar{g}_{ij}^{z}}}$: Correlation between vectors comprised of $\alpha^z$ and $\bar{g}_{ij}^{z}$ for $z = 1$ to $z = n$

$\rho_{\vec{\delta},\overrightarrow{\bar{g}_{ij}^{z}}}$: Correlation between vectors comprised of $\delta^z$ and $\bar{g}_{ij}^{z}$ for $z = 1$ to $z = n$

$\rho_{\vec{\alpha}^2,\overrightarrow{var}(g_{ij}^{z})}$ : Correlation between vectors comprised of $(\alpha^z)^2$ and $var\left(g_{ij}^{z}\right)$ for $z = 1$ to $z = n$

$\rho_{\vec{\delta}^2,\overrightarrow{var}(g_{ij}^{z})}$ : Correlation between vectors comprised of $(\delta^z)^2$ and $var\left(g_{ij}^{z}\right)$ for $z = 1$ to $z = n$

**A7. $\lambda$ is the Direct-Nurture Heritability Ratio**

$$\rho_{\vec{\alpha},\overrightarrow{\overline{g_{ij}^{z}}}} = \rho_{\vec{\delta},\overrightarrow{\overline{g_{ij}^{z}}}}$$

$$\frac{cov\left(\vec{\alpha},\overrightarrow{\overline{g_{ij}^{z}}}\right)}{var(\vec{\alpha})^{\frac{1}{2}}var\left(\overrightarrow{\overline{g_{ij}^{z}}}\right)^{\frac{1}{2}}} = \frac{cov\left(\vec{\delta},\overrightarrow{\overline{g_{ij}^{z}}}\right)}{var\left(\vec{\delta}\right)^{\frac{1}{2}}var\left(\overrightarrow{\overline{g_{ij}^{z}}}\right)^{\frac{1}{2}}}$$

$$\frac{cov\left(\vec{\alpha},\overrightarrow{\overline{g_{ij}^{z}}}\right)}{var(\vec{\alpha})^{\frac{1}{2}}} = \frac{cov\left(\vec{\delta},\overrightarrow{\overline{g_{ij}^{z}}}\right)}{var\left(\vec{\delta}\right)^{\frac{1}{2}}}$$

$$\frac{var\left(\vec{\delta}\right)^{\frac{1}{2}}}{var(\vec{\alpha})^{\frac{1}{2}}}cov\left(\vec{\alpha},\overrightarrow{\overline{g_{ij}^{z}}}\right) = cov\left(\vec{\delta},\overrightarrow{\overline{g_{ij}^{z}}}\right)$$

$$\frac{var\left(\vec{\delta}\right)^{\frac{1}{2}}}{var(\vec{\alpha})^{\frac{1}{2}}}[\sum_{z=1}^{n}(\alpha^{z}\overline{g_{ij}^{z}}\ ) - \bar{\alpha}\ \overline{\overline{g_{ij}^{z}}}] = \sum_{z=1}^{n}(\delta^{z}\bar{g}_{ij}^{z}) - \bar{\delta}\overline{\overline{g_{ij}^{z}}}$$

$$\frac{var\left(\vec{\delta}\right)^{\frac{1}{2}}}{var(\vec{\alpha})^{\frac{1}{2}}}[\sum_{z=1}^{n}(\alpha^{z}\overline{g_{ij}^{z}}) - 0] = \sum_{z=1}^{n}(\delta^{z}\overline{g_{ij}^{z}}) - 0$$

$$\frac{var\left(\vec{\delta}\right)^{\frac{1}{2}}}{var(\vec{\alpha})^{\frac{1}{2}}} = \frac{\sum_{z=1}^{n}(\delta^{z}\overline{g_{ij}^{z}})}{\sum_{z=1}^{n}(\alpha^{z}\overline{g_{ij}^{z}})}$$

$$\mathrm{E}\left[\frac{var\left(\vec{\delta}\right)^{\frac{1}{2}}}{var(\vec{\alpha})^{\frac{1}{2}}}\right] = \mathrm{E}\left[\frac{\sum_{z=1}^{n}(\delta^{z}\overline{g_{ij}^{z}})}{\sum_{z=1}^{n}(\alpha^{z}\overline{g_{ij}^{z}})}\right]$$

$$\frac{\left(\frac{\lambda^2\sigma}{4}\right)^{\frac{1}{2}}}{(\sigma)^{\frac{1}{2}}} = \mathrm{E}\left[\frac{\sum_{z=1}^{n}(\delta^{z}\overline{g_{ij}^{z}})}{\sum_{z=1}^{n}(\alpha^{z}\overline{g_{ij}^{z}})}\right]$$

$$\lambda = \mathrm{E}\left[\frac{2\sum_{z=1}^{n}(\delta^{z}\overline{g_{ij}^{z}})}{\sum_{z=1}^{n}(\alpha^{z}\overline{g_{ij}^{z}})}\right]$$

**A8. Variance of $\boldsymbol{PGS_{ij}^{\mathrm{N}}}$ is $\boldsymbol{\frac{\lambda^2}{4}}$**

$$\rho_{\vec{\alpha}^2,\overrightarrow{var}\left(g_{ij}^z\right))} = \rho_{\vec{\delta}^2,\overrightarrow{var}\left(g_{ij}^z\right)}$$

$$\frac{cov\left(\vec{\alpha}^2,\overrightarrow{var}\left(g_{ij}^z\right)\right)}{var(\vec{\alpha}^2)^{\frac{1}{2}}\overrightarrow{var}\left(g_{ij}^z\right)^{\frac{1}{2}}} = \frac{cov\left(\vec{\delta}^2,\overrightarrow{var}\left(g_{ij}^z\right)\right)}{var\left(\vec{\delta}^2\right)^{\frac{1}{2}}\overrightarrow{var}\left(g_{ij}^z\right)^{\frac{1}{2}}}$$

$$\frac{cov\left(\vec{\alpha}^2,\overrightarrow{var}\left(g_{ij}^z\right)\right)}{var(\vec{\alpha}^2)^{\frac{1}{2}}} = \frac{cov\left(\vec{\delta}^2,\overrightarrow{var}\left(g_{ij}^z\right)\right)}{var\left(\vec{\delta}^2\right)^{\frac{1}{2}}}$$

$$\frac{var\left(\vec{\delta}^2\right)^{\frac{1}{2}}}{var(\vec{\alpha}^2)^{\frac{1}{2}}} cov\left(\vec{\alpha}^2,\overrightarrow{var}\left(g_{ij}^z\right)\right) = cov\left(\vec{\delta}^2,\overrightarrow{var}\left(g_{ij}^z\right)\right)$$

$$\frac{var\left(\vec{\delta}^2\right)^{\frac{1}{2}}}{var(\vec{\alpha}^2)^{\frac{1}{2}}} \sum_{z=1}^{n} (\alpha^z)^2\, var(g_{ij}^z) - (\bar{\alpha}^z)^2 \overline{var}(g_{ij}^z) = \sum_{z=1}^{n} (\delta^z)^2\, var(g_{ij}^z) - (\delta^z)^2 \overline{var}(g_{ij}^z)$$

$$\frac{var\left(\vec{\delta}^2\right)^{\frac{1}{2}}}{var(\vec{\alpha}^2)^{\frac{1}{2}}} \sum_{z=1}^{n} (\alpha^z)^2\, var(g_{ij}^z) - 0 = \sum_{z=1}^{n} (\delta^z)^2\, var(g_{ij}^z) - 0$$

$$\frac{var\left(\vec{\delta}^2\right)^{\frac{1}{2}}}{var(\vec{\alpha}^2)^{\frac{1}{2}}}\sum_{z=1}^{n}(\alpha^z)^2\, var(g_{ij}^z) = \sum_{z=1}^{n}(\delta^z)^2\, var(g_{ij}^z)$$

$$\frac{var\left(\vec{\delta}^2\right)^{\frac{1}{2}}}{var(\vec{\alpha}^2)^{\frac{1}{2}}}\sum_{z=1}^{n} var(\alpha^z g_{ij}^z) = \sum_{z=1}^{n} var(\delta^z g_{ij}^z)$$

$$\frac{var\left(\vec{\delta}^2\right)^{\frac{1}{2}}}{var(\vec{\alpha}^2)^{\frac{1}{2}}} var(PGS_{ij}^{\mathrm{D}}) = var(PGS_{ij}^{\mathrm{N}})$$

$$\frac{var\left(\vec{\delta}^2\right)^{\frac{1}{2}}}{var(\vec{\alpha}^2)^{\frac{1}{2}}} = var(PGS_{ij}^{\mathrm{N}})$$

$$\frac{\left(\frac{\lambda^4\sigma^4}{16}\right)^{\frac{1}{2}}}{(\sigma^4)^{\frac{1}{2}}} = \mathrm{E}\left[var\left(PGS_{ij}^{\mathrm{N}}\right)\right]$$

$$\mathrm{E}\left[var\left(PGS_{ij}^{\mathrm{N}}\right)\right] = \frac{\lambda^2}{4}$$

**A9. Variance of $\boldsymbol{PGS_j^{\mathrm{N}}}$ is $\boldsymbol{\frac{\lambda^2}{2}}$**

$$var(PGS_j^{\mathrm{N}}) = var\left(PGS_j^{\mathrm{N_m}} + PGS_j^{\mathrm{N_p}}\right)$$

$$var(PGS_j^{\mathrm{N}}) = var(PGS_j^{\mathrm{N_m}}) + var\left(PGS_j^{\mathrm{N_p}}\right) + 2cov(PGS_j^{\mathrm{N_m}}, PGS_j^{\mathrm{N_p}})$$

$$\mathrm{E}[var(PGS_j^{\mathrm{N}})] = \mathrm{E}\left[var(PGS_{ij}^{\mathrm{N}}) + var(PGS_{ij}^{\mathrm{N}}) + 2cov(PGS_j^{\mathrm{N_m}}, PGS_j^{\mathrm{N_p}})\right]$$

$$\mathrm{E}[var(PGS_j^{\mathrm{N}})] = \frac{\lambda^2}{4} + \frac{\lambda^2}{4} = 0$$

$$\mathrm{E}[var(PGS_j^{\mathrm{N}})] = \frac{\lambda^2}{2}$$

**A10. Between Family Analyses**

$$\mathrm{E}[\hat{\psi}_1] = \frac{cov\left(\widehat{PGS'}^{\mathrm{D}}_{ij}, Y_{ij}\right)}{var\left(\widehat{PGS'}^{\mathrm{D}}_{ij}\right)}$$

$$\mathrm{E}[\hat{\psi}_1] = \frac{cov\left(\widehat{PGS'}^{\mathrm{D}}_{ij}, \widehat{PGS'}^{\mathrm{D}}_{ij}\right)\beta_1 + cov\left(\widehat{PGS'}^{\mathrm{D}}_{ij}, \widehat{PGS'}^{\mathrm{D}}_{ij}\right)\beta_2}{var\left(\widehat{PGS'}^{\mathrm{D}}_{ij}\right)}$$

$$\mathrm{E}[\hat{\psi}_1] = \frac{cov\left(\widehat{PGS'}^{\mathrm{D}}_{ij}, \widehat{PGS'}^{\mathrm{D}}_{ij}\right) + cov\left(\widehat{PGS'}^{\mathrm{D}}_{ij}, \widehat{PGS'}^{\mathrm{D}}_{ij}\right)}{var\left(\widehat{PGS'}^{\mathrm{D}}_{ij}\right)}$$

$$\mathrm{E}[\hat{\psi}_1] = cov\left(\frac{\widehat{PGS'}^{\mathrm{D}}_{ij} - \widehat{PGS'}^{\mathrm{D}}_{ij}}{var\left(\widehat{PGS'}^{\mathrm{D}}_{ij}\right)^{\frac{1}{2}}}, PGS^{\mathrm{D}}_{ij}\right) + cov\left(\frac{\widehat{PGS'}^{\mathrm{D}}_{ij} - \overline{\widehat{PGS'}^{\mathrm{D}}_{ij}}}{var\left(\widehat{PGS'}^{\mathrm{D}}_{ij}\right)^{\frac{1}{2}}}, PGS^{\mathrm{N}}_{j}\right)$$

$$\mathrm{E}[\hat{\psi}_1] = \frac{cov\left(\widehat{PGS'}^{\mathrm{D}}_{ij}, PGS^{\mathrm{D}}_{ij}\right) + cov\left(\widehat{PGS'}^{\mathrm{D}}_{ij}, PGS^{\mathrm{N}}_{j}\right)}{var\left(\widehat{PGS'}^{\mathrm{D}}_{ij}\right)^{\frac{1}{2}}}$$

$$\mathrm{E}[\hat{\psi}_1] = \frac{cov\left(PGS^{\mathrm{D}}_{ij} + PGS^{\mathrm{N}}_{ij}, PGS^{\mathrm{D}}_{ij}\right) + cov\left(PGS^{\mathrm{D}}_{ij} + PGS^{\mathrm{N}}_{ij}, PGS^{\mathrm{N}}_{j}\right)}{var\left(PGS^{\mathrm{D}}_{ij} + PGS^{\mathrm{N}}_{ij}\right)^{\frac{1}{2}}}$$

$$\mathrm{E}[\hat{\psi}_1] = \frac{cov\left(PGS^{\mathrm{D}}_{ij}, PGS^{\mathrm{D}}_{ij}\right) + cov\left(PGS^{\mathrm{N}}_{ij}, PGS^{\mathrm{D}}_{ij}\right) + cov\left(PGS^{\mathrm{D}}_{ij}, PGS^{\mathrm{N}}_{j}\right) + cov\left(PGS^{\mathrm{N}}_{ij}, PGS^{\mathrm{N}}_{j}\right)}{var\left(PGS^{\mathrm{D}}_{ij} + PGS^{\mathrm{N}}_{ij}\right)^{\frac{1}{2}}}$$

$$\mathrm{E}\left[\hat{\psi}_1\right] = \frac{1 + \frac{\lambda\rho_g}{2} + \frac{\lambda\rho_g}{2} + \frac{\lambda^2}{4}}{\left(1 + \lambda\rho_g + \frac{\lambda^2}{4}\right)^{\frac{1}{2}}}$$

$$\mathrm{E}\left[\hat{\psi}_1\right] = \frac{1 + \lambda\rho_g + \frac{\lambda^2}{4}}{\left(1 + \lambda\rho_g + \frac{\lambda^2}{4}\right)^{\frac{1}{2}}}$$

$$\mathrm{E}\left[\hat{\psi}_1\right] = \sqrt{1 + \lambda\rho_g + \frac{\lambda^2}{4}}$$

**A11. Within Family Analyses**

$$\mathrm{E}[\hat{\pi}_1] = \frac{cov\left(\Delta_0^1 \widehat{PGS}'^{\mathrm{D}}_{ij}, \Delta_0^1 Y_{ij}\right)}{var\left(\Delta_0^1 \widehat{PGS}'^{\mathrm{D}}_{ij}\right)}$$

$$\mathrm{E}[\hat{\pi}_1] = \frac{cov\left(\Delta_0^1 \widehat{PGS}'^{\mathrm{D}}_{ij}, \Delta_0^1 PGS^{\mathrm{D}}_{ij}\right)\beta_1 + cov\left(\Delta_0^1 \widehat{PGS}'^{\mathrm{D}}_{ij}, \Delta_0^1 PGS^{\mathrm{N}}_{j}\right)\beta_2}{var\left(\Delta_0^1 \widehat{PGS}'^{\mathrm{D}}_{ij}\right)}$$

$$\mathrm{E}[\hat{\pi}_1] = \frac{cov\left(\Delta_0^1 \widehat{PGS}'^{\mathrm{D}}_{ij}, \Delta_0^1 PGS^{\mathrm{D}}_{ij}\right) + cov\left(\Delta_0^1 \widehat{PGS}'^{\mathrm{D}}_{ij}, \Delta_0^1 PGS^{\mathrm{N}}_{j}\right)}{var\left(\Delta_0^1 \widehat{PGS}'^{\mathrm{D}}_{ij}\right)}$$

$$\mathrm{E}[\hat{\pi}_1] = \frac{cov\left(\Delta_0^1 \widehat{PGS}'^{\mathrm{D}}_{ij}, \Delta_0^1 PGS^{\mathrm{D}}_{ij}\right)}{var\left(\Delta_0^1 \widehat{PGS}'^{\mathrm{D}}_{ij}\right)}$$

$$\mathrm{E}[\hat{\pi}_1] = cov\left(\frac{\Delta_0^1 \widehat{PGS}^{\mathrm{D}}_{ij} - \Delta_0^1 \overline{\widehat{PGS}}^{\mathrm{D}}_{ij}}{var\left(\Delta_0^1 \widehat{PGS}^{\mathrm{D}}_{ij}\right)^{\frac{1}{2}}}, \Delta_0^1 PGS^{\mathrm{D}}_{ij}\right)$$

$$\mathrm{E}[\hat{\pi}_1] = \frac{cov\left(\Delta_0^1 \widehat{PGS}^{\mathrm{D}}_{ij}, \Delta_0^1 PGS^{\mathrm{D}}_{ij}\right)}{var\left(\Delta_0^1 \widehat{PGS}^{\mathrm{D}}_{ij}\right)^{\frac{1}{2}}}$$

$$\mathrm{E}[\hat{\pi}_1] = \frac{cov\left(\Delta_0^1 PGS^{\mathrm{D}}_{ij} + \Delta_0^1 PGS^{\mathrm{N}}_{ij}, \Delta_0^1 PGS^{\mathrm{D}}_{ij}\right)}{var\left(\Delta_0^1 PGS^{\mathrm{D}}_{ij} + \Delta_0^1 PGS^{\mathrm{N}}_{ij}\right)^{\frac{1}{2}}}$$

$$\mathrm{E}[\hat{\pi}_1] = \frac{cov\left(\Delta_0^1 PGS^{\mathrm{D}}_{ij}, \Delta_0^1 PGS^{\mathrm{D}}_{ij}\right) + cov\left(\Delta_0^1 PGS^{\mathrm{N}}_{ij}, \Delta_0^1 PGS^{\mathrm{D}}_{ij}\right)}{var\left(\Delta_0^1 PGS^{\mathrm{D}}_{ij} + \Delta_0^1 PGS^{\mathrm{N}}_{ij}\right)^{\frac{1}{2}}}$$

$$\mathrm{E}[\hat{\pi}_1] = \frac{var\left(\Delta_0^1 PGS_{ij}^{\mathrm{D}}\right)\beta_1 + cov\left(\Delta_0^1 PGS_{ij}^{\mathrm{N}}, \Delta_0^1 PGS_{ij}^{\mathrm{D}}\right)}{var\left(\Delta_0^1 PGS_{ij}^{\mathrm{D}} + \Delta_0^1 PGS_{ij}^{\mathrm{N}}\right)^{\frac{1}{2}}}$$

$$\mathrm{E}[\hat{\pi}_1] = \frac{1 + \frac{\lambda \rho_g}{2}}{\left(1 + \lambda \rho_g + \frac{\lambda^2}{4}\right)^{\frac{1}{2}}}$$

### A12. Empirical Results (Full Regression Table)

Table A1. The relationship between polygenic score and observed trait for six phenotypes in Add Health, within families and between families.

| | Years of Schooling | | Cognitive Ability | | CESD Depression Index | | Birth Weight | | Body Mass Index | | Height | |
|---|---|---|---|---|---|---|---|---|---|---|---|---|
| Family Fixed Effect | | X | | X | | X | | X | | X | | X |
| Educational Attainment PGS | 0.808** | 0.354** | | | | | | | | | | |
| | (0.0275) | (0.132) | | | | | | | | | | |
| Cognitive Ability PGS | | | 3.173** | 1.699** | | | | | | | | |
| | | | (0.158) | (0.629) | | | | | | | | |
| Depression PGS | | | | | 0.130** | 0.0556 | | | | | | |
| | | | | | (0.0132) | (0.0650) | | | | | | |
| Birth Weight PGS | | | | | | | 3.071** | 3.256* | | | | |
| | | | | | | | (0.317) | (1.360) | | | | |
| Body Mass Index PGS | | | | | | | | | 1.999** | 2.344** | | |
| | | | | | | | | | (0.0963) | (0.542) | | |
| Height PGS | | | | | | | | | | | 2.496** | 2.508** |
| | | | | | | | | | | | (0.0932) | (0.407) |
| r2 | 0.173 | 0.756 | 0.0836 | 0.736 | 0.0361 | 0.598 | 0.0353 | 0.816 | 0.0837 | 0.695 | 0.583 | 0.864 |
| N | 5323 | 734 | 5087 | 702 | 5323 | 734 | 4524 | 647 | 5269 | 727 | 5299 | 733 |

+ 0.10 * 0.05 ** 0.01 . All models control for sex, age, and the first 10 principal components of individual genotype. All model uses only individuals of European ancestry. Models without individual-fixed effects use a sample of unrelated individuals, whereas the family-fixed effect models use a sample of sibling pairs. The sample of unrelated individuals contains one randomly selected sibling from each sibling pair. All polygenic scores are both standardized within sample to be mean 0 and standard deviation 1. Cognitive ability is measured through the Peabody Picture Vocabulary Test during Wave 1 of Add Health, when respondents were approximately 16 years old. Birth weight is retrospectively reported by respondents' parents during Wave 1 of Add Health. Years of schooling, CESD depression index, body mass index, and height are measured during Wave 4 of Add Health, when respondents were approximately 28 years old. Height is reported in centimeters, birth weight is reported in ounces, and cognitive ability is reported in IQ score points. The CESD depression index is normalized to be mean 0 and standard deviation 1.
